# Acetylation of the yeast Hsp40 chaperone protein Ydj1 fine-tunes proteostasis and translational fidelity

**DOI:** 10.1101/2024.06.13.598777

**Authors:** Siddhi Omkar, Courtney Shrader, Joel R. Hoskins, Jake T. Kline, Megan M. Mitchem, Nitika, Luca Fornelli, Sue Wickner, Andrew W. Truman

## Abstract

Proteostasis, the maintenance of cellular protein balance, is essential for cell viability and is highly conserved across all organisms. Newly synthesized proteins, or “clients,” undergo sequential processing by Hsp40, Hsp70, and Hsp90 chaperones to achieve proper folding and functionality. Despite extensive characterization of post-translational modifications (PTMs) on Hsp70 and Hsp90, the modifications on Hsp40 remain less understood. This study aims to elucidate the role of lysine acetylation on the yeast Hsp40, Ydj1. By mutating acetylation sites on Ydj1’s J-domain to either abolish or mimic constitutive acetylation, we observed that preventing acetylation had no noticeable phenotypic impact, whereas acetyl-mimic mutants exhibited various defects indicative of impaired Ydj1 function. Proteomic analysis revealed several Ydj1 interactions affected by J-domain acetylation, notably with proteins involved in translation. Further investigation uncovered a novel role for Ydj1 acetylation in stabilizing ribosomal subunits and ensuring translational fidelity. Our data suggest that acetylation may facilitate the transfer of Ydj1 between Ssa1 and Hsp82. Collectively, this work highlights the critical role of Ydj1 acetylation in proteostasis and translational fidelity.

**Author Summary:** Cells require a suite of chaperone and co-chaperone proteins to maintain a healthy balance of functional proteins. A large number of modifications on chaperone and co-chaperone proteins have been identified, but their functional importance has not been fully explored. In this study, we identify acetylation sites on the yeast co-chaperone Ydj1 that impact its interactions with major chaperones and client proteins including those involved in protein synthesis. This work sheds light on how modifications on co-chaperones can also play an important role in the health of the proteome.

## Introduction

The highly dynamic and crowded cellular landscape presents a challenge for newly synthesized proteins that need to achieve and retain their active, native fold. Molecular chaperones such as Hsp70 and Hsp90 are directly involved in supporting protein function and assist in the assembly of multimeric complexes by binding “clients” as they are synthesized [1]. The activity and specificity of Hsp70 is directed by a suite of co-chaperone proteins primarily comprised of Hsp40s, also known as J-domain proteins (JDPs) and nucleotide exchange factors (NEFs) [2,3]. In budding yeast, 22 JDPs provide support for 11 Hsp70 paralogs [4]. The most highly expressed of these is Ydj1 at over 40,000 molecules per cell [4]. Perturbation of Ydj1 function results in a variety of phenotypes that include poor growth at room temperature and severe temperature sensitivity [5,6]. Cells lacking Ydj1 are also sensitive to a range of cell wall damaging agents such as caffeine, calcofluor white (CFW), and sodium dodecyl sulfate (SDS), probably as a result of the interplay between chaperones and the yeast cell integrity MAPK pathway [7,8]. Ydj1 and its homologs contain six major functional domains that contribute to their crucial role in cellular protein folding processes [2,9,10] (Fig. 1A and B). The N-terminally located domain (J-domain) is required for stimulation of the ATPase activity of Hsp70, facilitating the transfer of client proteins from Hsp40 to Hsp70 for further processing [11,12]. Studies of this region have identified a highly conserved sequence His-Pro-Asp (HPD) essential for Hsp70-Hsp40 interaction [13,14]. This domain is followed by a Glycine/Phenylalanine-rich region, which helps determine client specificity [15]. Following the Glycine/Phenylalanine-rich region, there are two carboxy-terminal domains (CTDs), which act as key client-binding domains [16]. The first of these domains (CTDI) contains a client-binding site. Within the Ydj1 CTDI is a zinc-finger-like region, which aids in the stabilization of client proteins [17]. The second CTD, CTDII, also binds clients and is paramount for J-protein self-regulation [12,17]. This regulatory role is made possible with help from the dimerization domain. Following the dimerization domain of Ydj1 is a C-terminal extension. While its function is currently undefined, a similar region in human DNAJA2 regulates self-association and chaperone activity [18].

**Figure 1.**
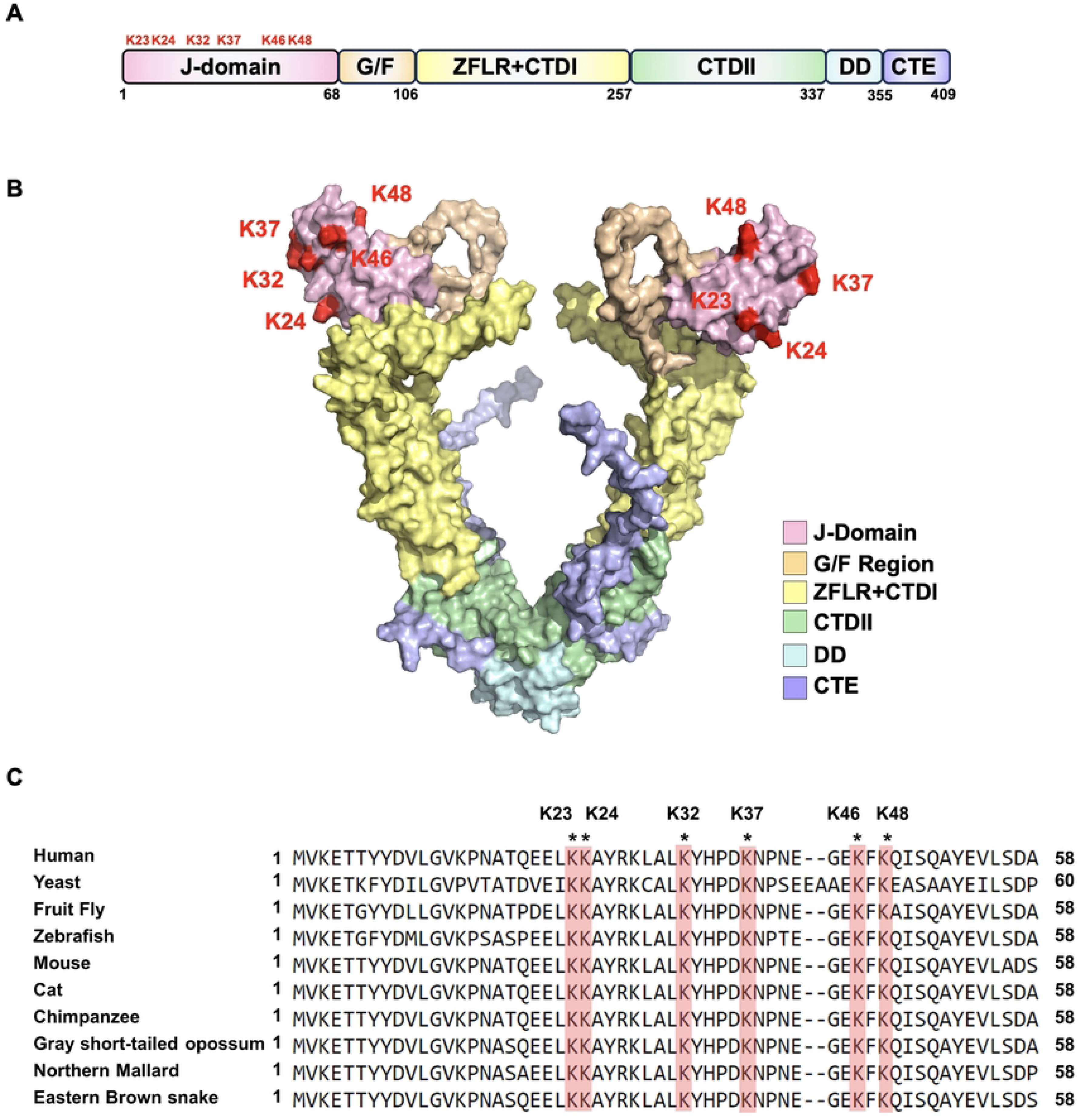
Yeast Ydj1 contains six acetylation sites in its J-domain. (A) Domain structure of yeast Ydj1. Sites of acetylation on the J-domain are highlighted in red. (B) Cartoon representation of Ydj1 with individual domains colored as indicated. Acetylated residues are labeled in red. The structure was generated by Alphafold2 and rendered in PyMOL. (C) Alignment of Ydj1 homologs showing the location and conservation of J-domain acetylated lysines.

Despite several decades of research, the only post-translational modification on Ydj1 that has been extensively studied is C-terminal farnesylation [19–21]. While Ydj1 is primarily cytosolic, farnesylation at cysteine 406 facilitates the localization of Ydj1 to the Endoplasmic Reticulum and the perinuclear membrane [21,22]. This modification is crucial for the regulation and activation of specific Hsp90 clients [21]. Recent studies have shown that in mammalian cells, Hsc70 and its co-chaperone DNAJA1 are deacetylated by HDAC6, a process required for the interaction between the two proteins [23]. In this study, we have explored the impact of Ydj1 J-domain acetylation on its function both in vivo and in vitro. Our findings suggest that Ydj1 acetylation plays a vital role in proteostasis and translational fidelity.

## Results

### Ydj1 acetylation impacts the yeast stress response

Global proteomic experiments have identified 12 acetylation sites on Ydj1 (GPMdb, https://gpmdb.thegpm.org/). To prioritize sites for further study, we considered their conservation throughout nature and their location on important functional domains. Six acetylation sites, K23, K24, K32, K37, K46, and K48 are found on the j-domain of Ydj1, a region essential for interaction with Ssa1 and Ydj1 function [3,13,19] (Figure 1A and B). Interestingly, these sites are highly conserved in eukaryotes, suggesting potential functional importance[19] (Figure 1C). To assess the influence of acetylation on yeast stress resistance, we constructed a yeast centromeric plasmid expressing Ydj1 with a C-terminal FLAG tag from a constitutive GPD promoter. We individually substituted each of the six J-domain lysines with either arginine (non-acetylatable) or glutamine (acetyl-mimic) in a manner similar to [24–27]. Additionally, we introduced combined substitutions of all six J-domain lysines to either arginine or glutamine (hereafter denoted as 6KR or 6KQ, respectively). The functionality of Ydj1 is crucial for the response to a range of cell stresses, including high temperature, DNA damage and agents that damage cell wall integrity [22]. To investigate the impact of acetylation on Ydj1 function, *ydj1*Δ cells were transformed with plasmids carrying an empty vector control or expressing wild-type Ydj1 or acetylation site mutants (R and Q versions of K23, K24, K32, K37, K46, K48). While cells lacking Ydj1 displayed increased sensitivity to all stress conditions compared to the wild-type, those expressing any of the six arginine substitutions grew comparably to wild-type, indicating that preventing acetylation of residues in the Ydj1 J-domain minimally affects yeast stress resistance. (Figure 2A). In contrast, substituting J-domain lysines with glutamine (KQ) had a more pronounced effect on Ydj1 function. Notably, cells expressing Ydj1 K23Q and K37Q failed to thrive at elevated temperatures. (Figure 2B). Interestingly, this effect was not universal across all tested stress conditions. Although K37Q displayed a partially caffeine-sensitive phenotype, it mirrored wild-type Ydj1 function under other stress conditions (Figure 2B). K23Q and K24Q cells grew at wild-type rates in response to caffeine, Calcofluor white (CFW), SDS, and UV treatment (Figure 2B). Mutation of all six sites to arginine (6KR) had no discernible impact on the cellular response to the tested stresses (Figure 2C). Conversely, the 6KQ mutation produced the most notable loss-of-function phenotype, exhibiting sensitivity to all stresses examined (Figure 2C).

**Figure 2.**
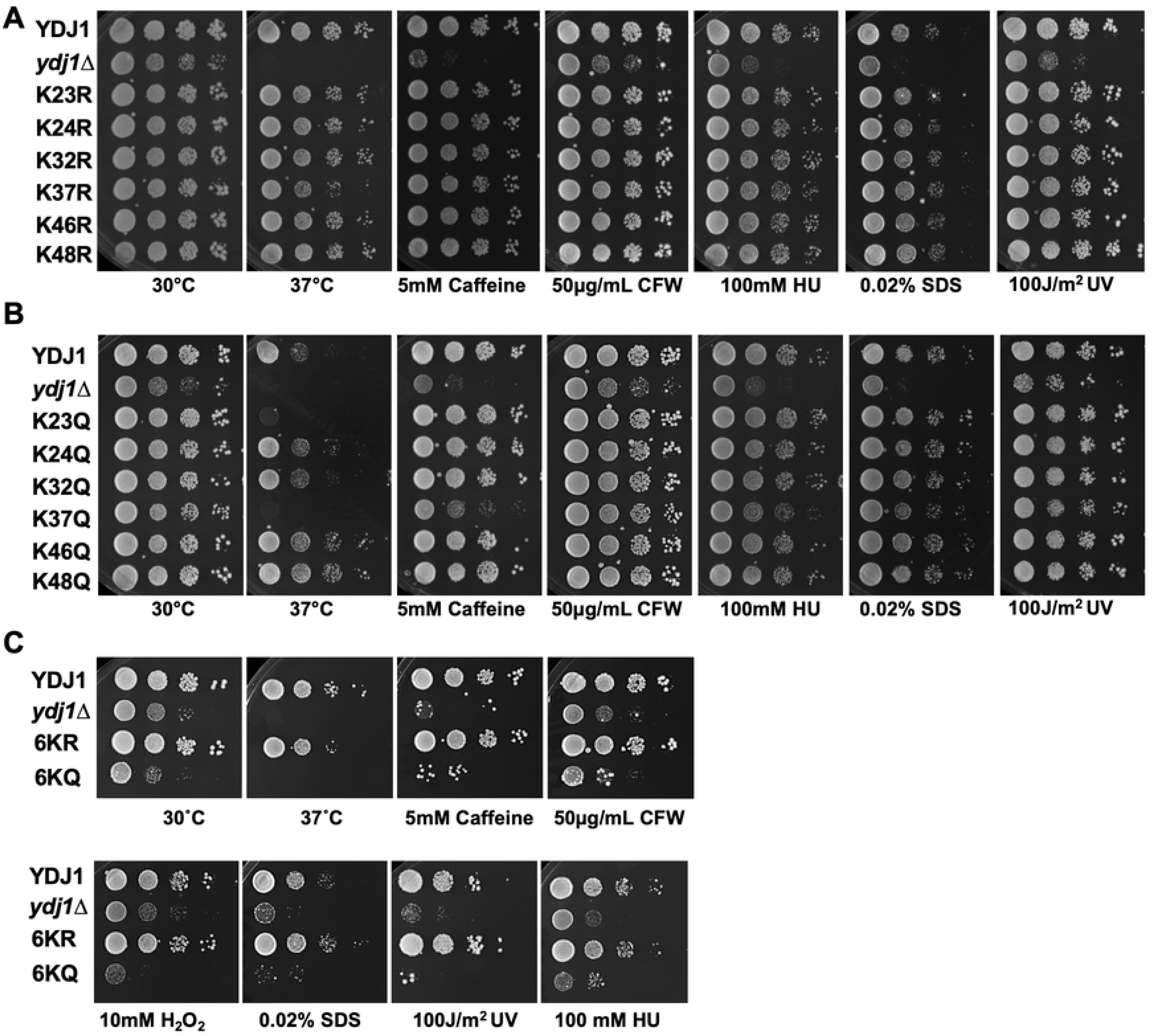
Ydj1 acetylation impacts yeast stress resistance. (A) Serial dilution of yeast expressing mutations that prevent acetylation of residues in the Ydj1 J-domain. WT YDJ1 and *ydj1*Δ cells transformed with either an empty control plasmid or a plasmid expressing a single Ydj1 KR mutant were grown to mid-log phase and then tenfold serially diluted onto YPD media containing the indicated stress agents. Plates were photographed after three days. (B) Serial dilution of yeast expressing mutations that mimic acetylation of residues in the Ydj1 J-domain. WT YDJ1 and *ydj1*Δ cells transformed with either a control plasmid or plasmid expressing a single Ydj1 KQ mutant were grown to mid-log phase and then tenfold serially diluted onto YPD media containing the indicated stress agents. Plates were photographed after three days. (C) Serial dilution of yeast expressing 6KR/6KQ mutations. WT YDJ1 and *ydj1*Δ cells transformed with either a control plasmid, Ydj1-6KQ, or Ydj1-6KR were grown to mid-log phase and then tenfold serially diluted onto YPD media containing the indicated stress agents. Plates were photographed after three days.

### Ydj1 acetylation alters Ydj1 abundance and mobility on SDS-PAGE

Acetylation can impact a range of protein properties, including protein stability [28]. Several of the Ydj1 KQ mutants were defective for growth under several conditions, suggesting that acetylation may alter Ydj1 stability. We analyzed the Ydj1 abundance in cells expressing Ydj1 wild-type and lysine substitutions from plasmids via Western blot analysis. The majority of the single KR mutants displayed levels of Ydj1 comparable to the wild-type except for K46R, which exhibited a subtle decrease in Ydj1 (Figure 3A). Similarly, the abundance of Ydj1 was not significantly impacted in the majority of single KQ mutants except for K37Q, which displayed a large increase in Ydj1 (Figure 3A). Interestingly, this was not observed in the 6KQ mutant, suggesting a potential cooperative effect between the six acetylation sites (Figure 3A). As seen previously, Ydj1 wild-type appeared as a doublet by Western blot analysis due to the presence of both unfarnesylated and farnesylated forms [29]. All of the Ydj1 mutants were also observed as doublet bands by Western blot analysis, indicating that farnesylation and J-domain acetylation are independently regulated processes (Figure 3A). Interestingly, we also noted an apparent increase in size of Ydj1 in K46Q, K48Q and 6KQ mutants (Figure 3A) Overall, these observations demonstrate that even a single K to R mutation can impact Ydj1 abundance and apparent size on SDS-PAGE. To determine whether this apparent increase in size and abundance was dependent on expression in yeast, we constructed plasmids carrying C-terminally His-tagged Ydj1 wild-type and mutants and expressed the proteins in *E. coli*. A single band for Ydj1 was observed after SDS-PAGE followed by Western blot analysis, presumable due to the lack of farnesylation machinery in bacteria. Interestingly, the mobility difference observed between the 6KQ and 6KR mutants was still present in the recombinant system (Figure 3B). A well-established consequence of purifying recombinant Ydj1 is the co-purification of a Ydj1 fragment lacking the J-domain (known as delta J). After purification of wild-type and mutant recombinant Ydj1 from bacteria, we analyzed Coomassie-stained SDS-PAGE gels for the presence of the delta J fragment. Delta J was present in all samples with a consistent apparent molecular weight, confirming that the size alterations seen in the full-length protein are due to the J-domain (Figure S1).

**Figure 3.**
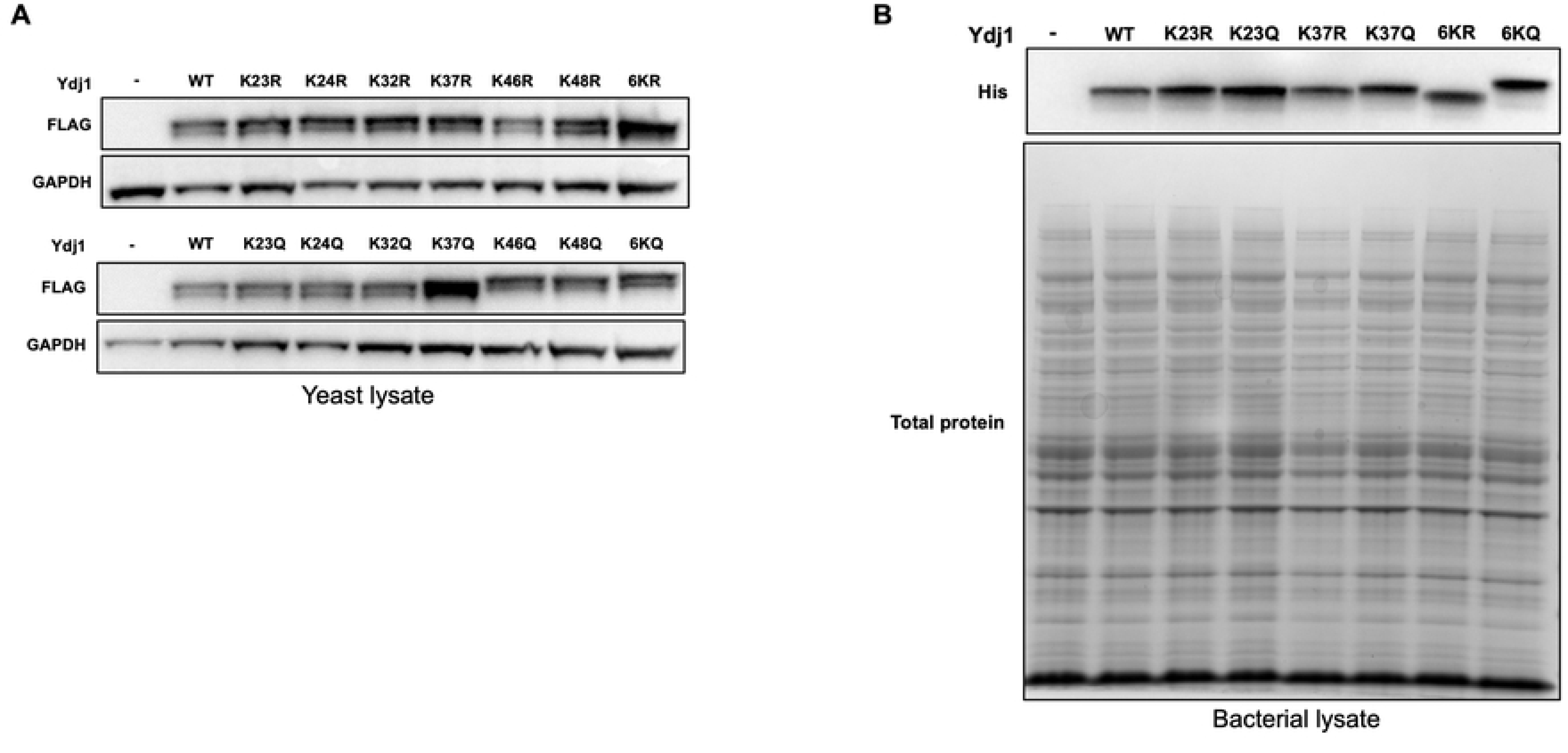
Impact of acetylation on steady-state levels and gel mobility of Ydj1. (A) Western blot analysis of FLAG-Ydj1 abundance from *ydj1*Δ yeast cells transformed with either a control plasmid or a plasmid expressing the indicated mutant. Lysates were probed with antisera to either FLAG or Pgk1 (loading control). (B) Western blot analysis of HIS-Ydj1 abundance from *E. coli* cells transformed with either a control plasmid or a plasmid expressing the indicated Ydj1 mutant. Lysates were probed with antisera to HIS and total protein was assessed by Ponceau staining of the membrane.

### J-domain acetylation impacts the global Ydj1 interactome

The interactions between chaperones and co-chaperones are modulated by cellular stress and the fine-tuning of stress response through post-translational modifications, also known as the chaperone code [30]. To determine whether Ydj1 protein interactions are modulated by J-domain acetylation, we quantitatively compared the interactomes of Ydj1-FLAG 6KQ and Ydj1-FLAG 6KR. Ydj1-FLAG complexes from cells expressing the 6KQ or 6KR mutants were isolated via FLAG-Dynabeads. These complexes were digested via trypsin, and peptides were comparatively analyzed using LC-MS (Figure 4A). Protein interactions of Ydj1 were normalized against the bait (Ydj1), and then the log2 interaction ratio (6KQ/6KR) was calculated. A change in the interaction between 6KQ and 6KR samples was considered significant if the normalized log2 (6KQ/6KR) value was >1 or <-1. The interactors were then sorted according to their Gene Ontology terms and were plotted with the y-value corresponding to a change in chaperone interaction (Figure 4B). Three hundred twenty-seven high-confidence Ydj1 interactors were identified, and 63% of these were unaffected by Ydj1 acetylation (Figure 4B). Consistent with the loss of Ydj1 function seen in the 6KQ mutant, 21% of Ydj1 interactors had a strong preference for the 6KR mutant, with 16% displaying an enhanced affinity for the 6KQ sample (Figure 4B and 4C). Gene Ontology (GO) analysis uncovered differences in the classes of proteins enriched for interaction between the 6KR and 6KQ samples (Figure 4C). For example, proteins involved in vesicle trafficking, chromatin regulation, mitochondrial function, and tRNA wobble modification tended to be enriched in 6KR complexes. In contrast, proteins involved in protein turnover, metabolism, and cell polarity were enriched in the 6KQ complexes (Figure 4C). Many of the Ydj1 interactions (38%) identified were proteins involved in protein translation. 13% of these showed a preference for the 6KQ mutant, while 10% of these showed a preference for the 6KR version (Figure 4B).

**Figure 4.**
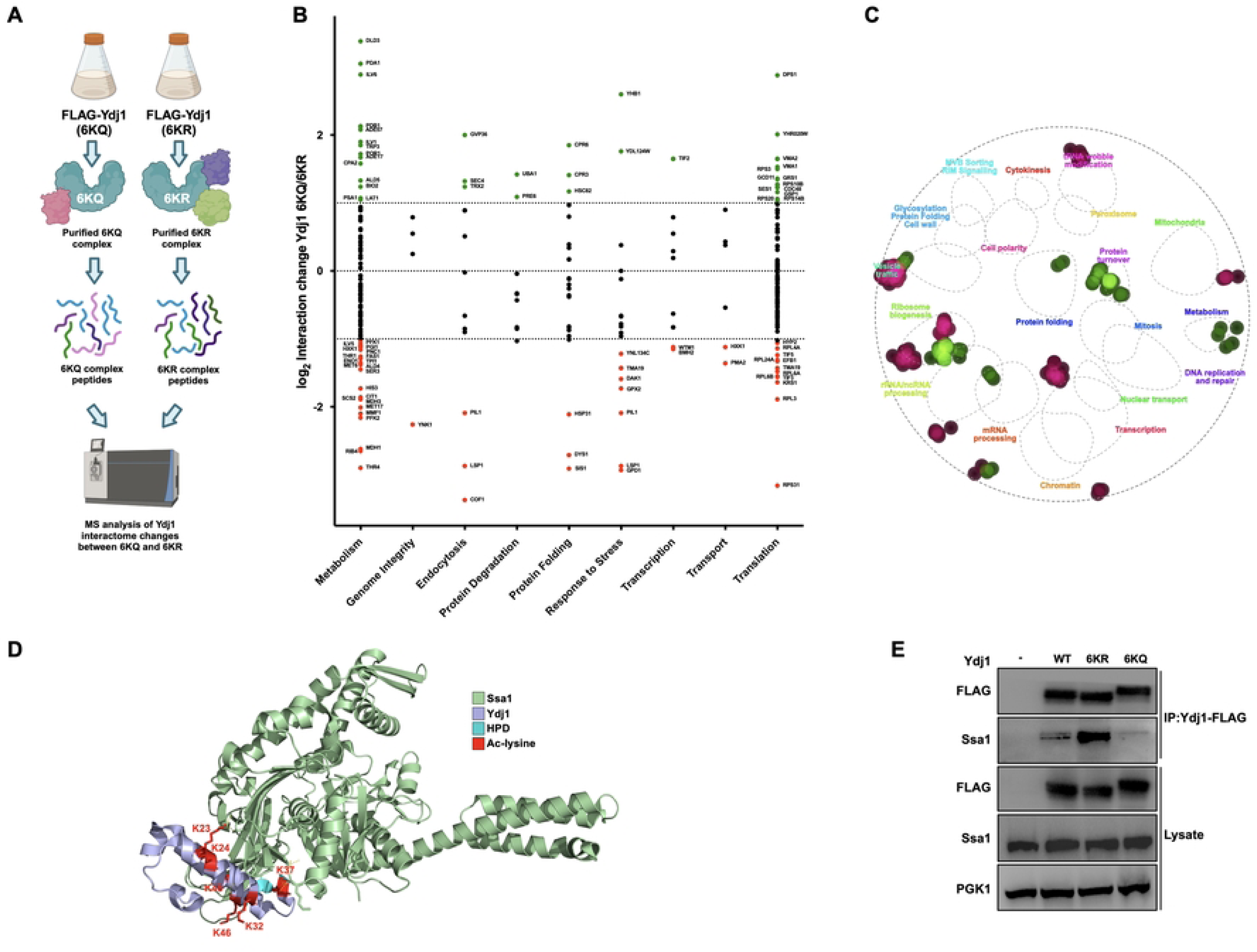
Ydj1 acetylation alters interactions with client proteins. (A) Experimental workflow of mass spectrometry analysis of Ydj1 complexes purified from yeast cells. *ydj1*Δ yeast cells expressing either FLAG-6KQ or 6KR variants of Ydj1 were grown to mid-log phase, and Ydj1 complexes were isolated via FLAG-dynabeads. 6KQ and 6KR complexes were digested into peptides via trypsin and were analyzed by mass spectrometry. (B) Comparative interactome analysis of Ydj1 6KQ and 6KR complexes. Interactors were organized into functional categories following GO terms and plotted against 6KQ/6KR interaction change (Log2 ratio). The dotted lines represent an interaction change of Log2 > 1 or Log2 < −1. Interactors are colored according to change in interaction as follows: green (significant increase), red (significant decrease), or black (no significant change) 6KQ/6KR. (C) GO analysis of Ydj1 interactors significantly increased (green) or decreased (red) upon Ydj1 acetylation (6KQ/6KR) using TheCellMap.org. (D) Acetylated lysine sites on Ydj1 are located on the interaction surface between Ydj1 and Ssa1. The Ydj1-Ssa1 complex was modeled was based on PDB entry 5NRO and rendered in PyMOL. The six acetylation sites are labeled in red. (E) Acetylation of Ydj1 disrupts interaction with Ssa1. Indicated Ydj1 complexes were isolated from yeast using FLAG-dynabeads, and interaction with Ssa1 was assessed via Western Blotting using antisera to FLAG and Ssa1. PGK1 was used to assess the equal loading of lysate samples.

Ydj1 mediates the majority of its functions through interactions with Hsp70 and Hsp90 chaperones as well as assorted co-chaperone proteins [31]. Modeling of the Ssa1-Ydj1 J-domain complex using the DnaK-J-domain crystal structure as a template (PBD 5NRO, reference) showed that all six Ydj1 acetylation sites are in close proximity to the interaction surface with Ssa1 (Figure 4D). The model predicted that two of these sites, K23 and K37, directly interact with the NBD of Ssa1 (Figure 4D). We considered the possibility that the altered interactome observed in Figure 4B may be a result of the altered Ydj1-Ssa1 association. In our interactome study, Ydj1 co-purified with Hsp70 paralogs Ssa1 and 2 and Hsp90 paralogs Hsc82 and Hsp82 (Figure 4B, Table S2). We also identified important members of the proteostasis network including Cpr6, Sis1, Ssb1/2, Sse1/2, Ssc1, Ssz1, Tsa1, Frp1, Pdi1, Hsp26, Hsp60, Hsp104 and Cpr3 (Figure 4B, Table S2). While the majority of Ydj1 chaperone and co-chaperone interactions were unchanged between the Ydj1 6KR and 6KQ interactomes, increased interaction was detected between Ydj1 and Hsc82, Cpr6 and Cpr3 in the 6KQ sample. In contrast, the interaction between Ydj1 and the related co-chaperone Sis1 was decreased in the 6KQ sample (Figure 4B).

Proteomic analysis of yeast Hsp70 Ssa chaperones is complicated due to their highly similar amino acid sequences [32]. We performed a direct immunoprecipitation experiment to clarify whether the Ydj1-Ssa1 interaction was truly perturbed by acetylation. Yeast expressing wild-type, 6KQ, or 6KR Ydj1-FLAG were grown to mid-log, the protein was extracted, and Ydj1 complexes were isolated by FLAG-Dynabeads. Immunoprecipitates were probed for the presence of Ssa1 using antisera specific to Ssa1 [33]). An increase in Ydj1-Ssa1 interaction was observed in the 6KR mutant, while almost no interaction was seen between the two proteins in the 6KQ mutant (Figure 4E). Overall, this data, together with the proteomic data (Figure 4A) suggests that acetylation may promote the transfer of Ydj1 from Ssa1 to Hsc82.

### J-domain acetylation of Ydj1 fine-tunes translational fidelity

Chaperones bind newly translated proteins and help them achieve their native, active state [34]. It is thus unsurprising that we identified numerous proteins involved in protein translation in our proteomics screen. In particular, Ydj1 co-purified with proteins involved in tRNA processing, ribosome structure, translational initiation, and elongation (Figure 5A). To understand if there was a spatial specificity for acetylation-dependent Ydj1 interactors, we mapped the identified Ydj1 interactors to the structure of the yeast ribosome (PDB: 4V7R) [35] (Figure 5B). Proteins clustered together in the ribosome’s small (40S) subunit tended to prefer interaction with the 6KQ mutant, whereas those in the large (60S) subunit decreased in interaction (Figure 5B). Changes in protein interactions in proteomic studies can be a result of either changes in the strength of association or altered abundance of the interactor. To determine whether ribosomal protein hits in our screen were caused by changes in protein abundance, we utilized the Dharmacon Yeast ZZ-tag ORF collection (https://horizondiscovery.com/en/non-mammalian-research-tools/products/yeast-orf-collection). These plasmids express ZZ-HA-HIS-tagged yeast ORFs from the inducible galactose promoter. We constitutively expressed three ribosomal proteins (Rps3, Rps0 and Gcd11) in wild-type and Ydj1 acetylation mutant cells (Figure 5C). The stability of Rps0 was dramatically reduced in cells lacking Ydj1 and moderately reduced in 6KQ cells (Figure 5C). Excitingly, the stability of Rps3 and Gcd11 was also clearly dependent on Ydj1 function, with lowered expression of both proteins in cells either lacking Ydj1 or expressing the KQ mutant (Figure 5C). Given these acetylation-dependent changes in the ribosomal proteins, we considered the possibility that acetylation may alter translational fidelity. We assessed translational fidelity in WT and Ydj1 mutant cells using a well-established dual luciferase assay. In this system, *Renilla* luciferase (Rluc) and firefly luciferase (Fluc) are under the control of separate constitutive promoters [36] (Figure 6D). The Fluc mRNA has a non-traditional start codon (UUG) which is only used when there is a loss of translational fidelity [36]. The Rluc mRNA has a classic AUG start codon and therefore acts as a control for alterations in overall translation. Using this system, we observed a loss of translational fidelity in cells lacking Ydj1, a phenotype that was recapitulated in KQ, but not KR cells (Figure 5E).

**Figure 5.**
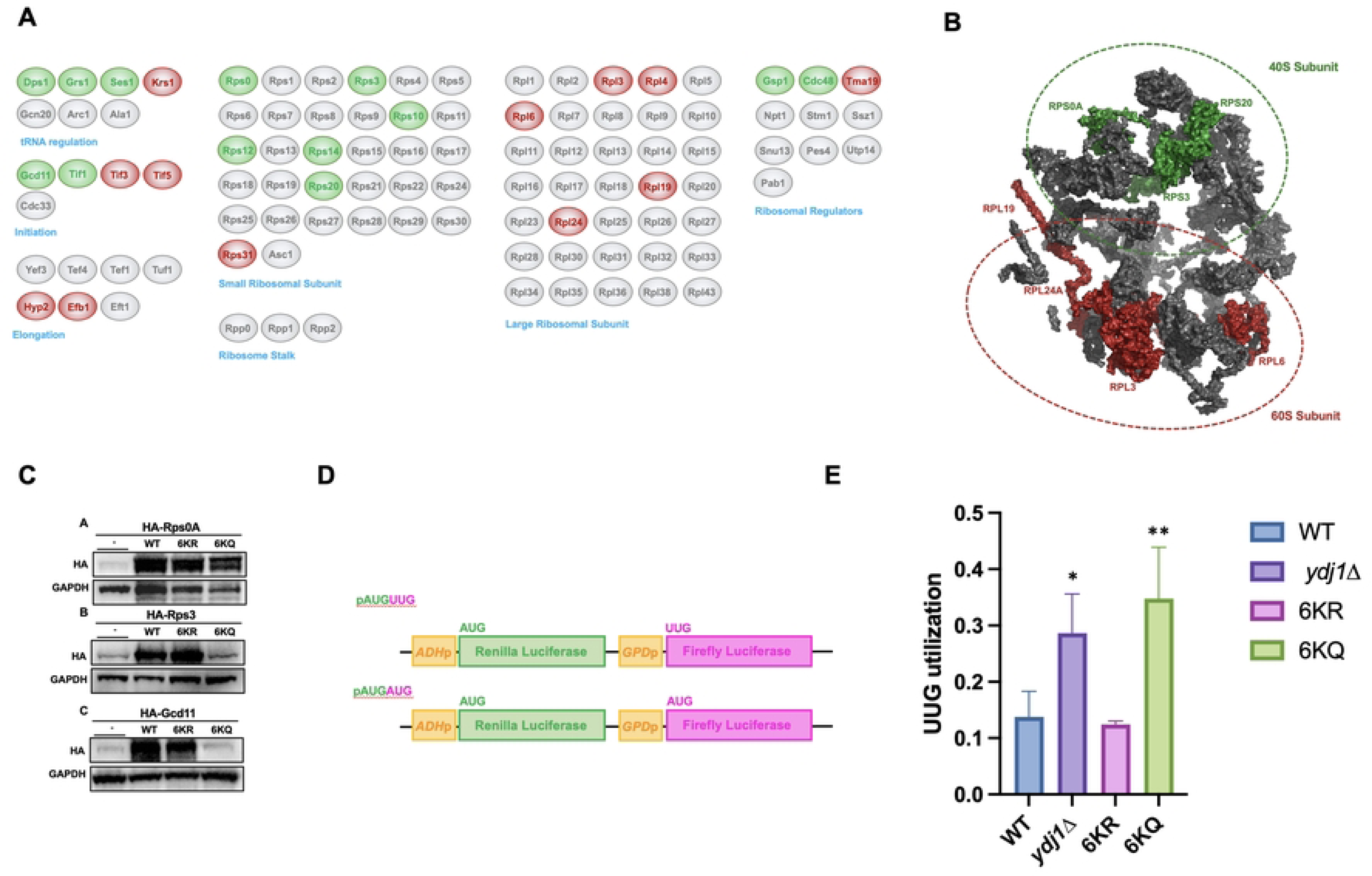
Ydj1 acetylation impacts multiple aspects of translation. (A) Interaction of Ydj1 with machinery involved in protein translation are impacted by Ydj1 acetylation. Interactors from the previous MS experiment were grouped by function and colored based on preference for either KR or KQ variant of Ydj1. Green, interactor has a preference for binding the KQ version: Grey, interactions are unaffected by Ydj1 acetylation; Red, interactor has a preference for the KR variant of Ydj1. (B) Ribosomal proteins displaying acetylation-dependent interaction with Ydj1 were mapped to the structure of the yeast ribosome (PDB: 4V7R). Interactors in green are those that increased in interaction, while those in red are interactors that showed a decrease in interaction (6KQ/6KR). (C) Ydj1 function impacts the stability of selected proteins involved in translation. *ydj1*Δ cells expressing plasmids encoding Ydj1 J-domain mutant proteins were transformed with plasmids for the expression of galactose-driven HA-tagged ribosomal proteins. Abundance of ribosomal proteins was assessed by Western blot analysis using antisera to HA and GAPDH. (D) Schematic of the translational fidelity assay used. (E) The fidelity of translation initiation in Ydj1 acetylation mutants was determined using the UUG/AUG dual luciferase assay described in [36]. The data shown are the mean and standard deviation of three biological replicates. Statistical significance was calculated via ANOVA. *, P≤ 0.05; **, P≤ 0.01.

**Figure 6.**
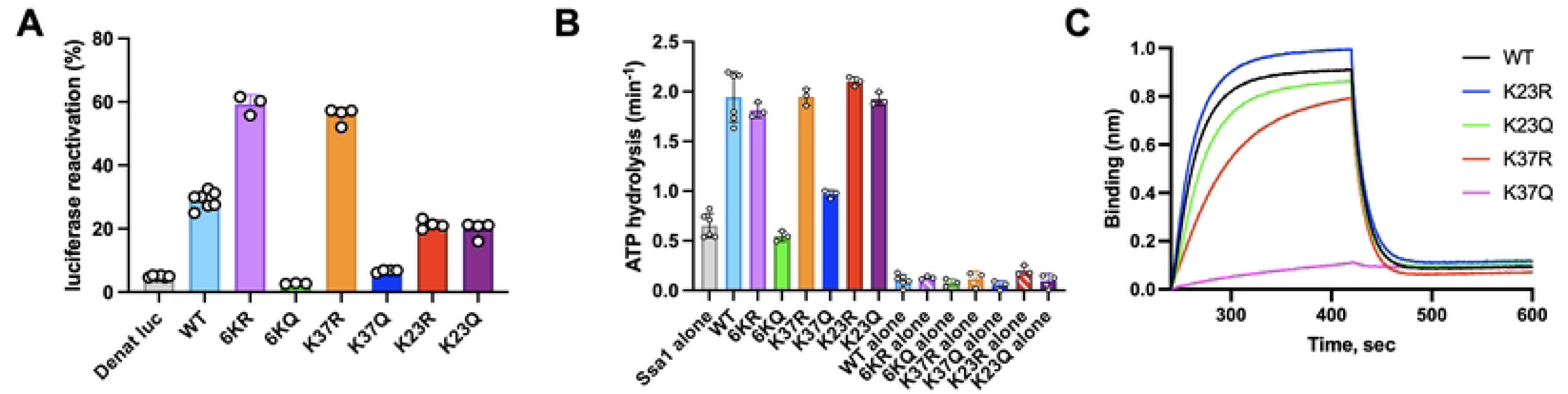
Acetylation impacts *in vitro* function of Ydj1. (A) Reactivation of chemically denatured luciferase was monitored over time in the presence of 3 μM Ssa1 and 0.3 μM Ydj1 wild-type or mutant as described in Materials and Methods. The percentage of luciferase reactivation after 40 min relative to a non-denatured luciferase control is shown. (B) The rate of Ssa1 ATP hydrolysis was determined in the absence and presence of Ydj1 wild-type or Ydj1 mutants 6KR or 6KQ as described in Materials and Methods. Background ATPase from the individual Ydj1 proteins is also shown. (C) Ssa1 ATPase as described in (B) using Ydj1 wild-type and mutants K23R, K23Q, K37R and K37Q. (D) The interaction of Ssa1 with Ydj1 wild-type or Ydj1 mutant proteins was analyzed using BioLayer Interferometry as described in Materials and Methods. Biotinylated Ydj1 wild-type or mutant was loaded onto streptavidin biosensors to equivalent levels and the association and dissociation of Ssa1 was monitored. The mean from three independent experiments is shown. In A, B and C results are shown as means ± SD.

### The *in vitro* function of Ydj1 is directed through acetylation

To understand the fundamental reason for the loss of Ydj1 function in the KQ mutants, we performed a series of *in vitro* assays using Ydj1 wild-type and mutant proteins that were expressed in *E. coli* and purified. Since protein acetylation does not occur in *E. coli*, the purified proteins all lacked the natural acetylation seen in proteins expressed in yeast. Thus, the wild-type Ydj1 would be expected to mimic the 6KR mutant. Initially, we measured the ability of wild-type Ydj1 and the Ydj1 arginine and glutamine substitution mutants to assist Ssa1 in the refolding of a model substrate, denatured luciferase (Figure 6A). The Ydj1 6KQ and K37Q mutants were defective in luciferase reactivation, whereas the 6KQ and K37Q mutants exhibited ∼two-fold enhanced ability to refold luciferase compared to wild-type (Figure 6A). This observation suggests acetylation of the Ydj1 J-domain, and in particular residue K37, is important for the function of Ydj1 in protein reactivation. The increased level of reactivation seen by 6KR and K37R suggests that a lysine to arginine substitution in residue 37 results in a protein more active in molecular chaperone activity with Ssa1. K23R and K23Q mutants were indistinguishable from wild-type, suggesting that acetylation of K23 does not impact the function of Ydj1 with Ssa1.

Since the ability of Ydj1 to assist in client refolding is dependent on its collaboration with Ssa1, we hypothesized that the defect in luciferase refolding may be a result of impaired stimulation of ATP hydrolysis of Ssa1. When wild-type Ydj1, 6KR, and 6KQ were tested for their ability to stimulate ATP hydrolysis by Ssa1, we observed that the Ydj1 6KQ mutant was defective (Fig. 6B). 6KQ stimulated Ssa1 ATPase to the same extent as wild-type Ydj1. To determine the impact of individual Ydj1 lysine substitutions on Ssa1 activity, we monitored the effect of K23Q, K23R, K37Q and K37R on ATP hydrolysis by Ssa1. The K37Q mutant was unable to stimulate Ssa1 activity. The K37R mutant as well as either K23Q or K23R) did not affect ATP hydrolysis (Figure 6C).

Since Ydj1 K37 is located next to the conserved HPD motif, which is known to interact with Hsp70s [14], and both K23 and K37 lie on the interface between Ydj1 and Ssa1 (Fig. 4D), we next tested the possibility that these residues participate in the interaction between Ydj1 and Ssa1. Binding between biotinylated Ssa1 and Ydj1 wild-type and mutants was monitored by BLI (Fig. 6D). Our results showed that Ydj1 K37Q was very defective in its ability to interact with Ssa1, while K37R was partially defective. K23R and K23Q were unchanged in their ability to bind Ssa1. Altogether these in vitro results confirm the in vivo results demonstrating that acetylation of K37 renders Ydj1 inactive and further show that acetylation inhibits the cochaperone activity of Ydj1 because it is unable to interact with Ssa1.

## Discussion

Over the past decade, research has unveiled a “Chaperone Code”—a system of post-translational modifications (PTMs) that regulate molecular chaperone function [30,37–39]. Despite the identification of thirty PTM sites on Ydj1, their specific roles and regulatory mechanisms remain largely unknown. Lysine acetylation is a rapidly reversible PTM tightly controlled through lysine acetyltransferases (KATs) and lysine deacetylases (KDACs) [23,40]. In *Saccharomyces cerevisiae*, around 4000 lysine acetylation sites have been uncovered on proteins involved in DNA repair, chromatin remodeling, cellular metabolism, protein folding, and transcription (via histone acetylation/deacetylation) [40,41]. In this study, we set out to determine the relevance of Ydj1 acetylation. We identified six acetylation sites that were highly conserved and present in the J-domain of Ydj1, a region critical for Ydj1 function. Mutation of these sites to either prevent or mimic acetylation at these sites produced an interesting spectrum of phenotypes. Surprisingly, prevention of acetylation produced no detectable effect. In contrast, K23Q, K37Q, and 6KQ showed a complete loss of growth at high temperatures. The J-domain of Ydj1 forms an interaction surface that allows the HPD region to bind to and stimulate the ATPase activity of Ssa1, thereby closing Ssa1’s lid and trapping the client protein [2]. This activity is highly regulated, though there is still some debate as to the current mechanisms behind this. Importantly, K23 and K37 lie directly on the interaction surface and participate in hydrogen bonding between Ydj1 and Ssa1. The other four lysines studied here are extremely close to the interaction surface as well. Our *in vivo* and *in vitro* data clearly demonstrate that interaction between Ssa1 and Ydj1 is disrupted by Ydj1 acetylation at K23 and K37. In accordance with our cellular data, Ydj1 K23Q and K37Q have a reduced ability to bind Ssa1 and stimulate its ATPase activity. This is also apparent in the inability of these mutants to promote the correct folding of the model luciferase substrate. Overall, the main phenotypic defects seen in the KQ mutants are likely a result of the loss of ability to bind Ssa1 and stimulate the Ssa1 ATPase activity.

Given the potential disruption of Ydj1-Ssa1 interaction by acetylation, we anticipated that the acetylation of Ydj1 would have far-reaching effects on its global protein interactions. In our MS experiment, we identified 327 interactors of Ydj1, 68% of which were unaffected by the acetylation status of Ydj1, and 16% of Ydj1 interactions were enriched in the KQ, suggesting that acetylation subtly fine-tunes Ydj1 interactions. Despite structural evidence as well as *in vitro* and *in vivo* assays showing a clear impact of acetylation on the Ssa1-Ydj1 interaction, this loss was not recapitulated in our mass spectrometry analysis. Our explanation for this discrepancy lies in the challenges of distinguishing the highly similar Ssa isoforms Ssa1, 2, 3, and 4 by mass spectrometry. To bypass this issue, we performed a co-immunoprecipitation/Western Blot analysis of the Ydj1-Ssa1 interaction using antisera specific to the Ssa1 isoform. Consistent with our original hypothesis, acetylation of Ydj1 indeed disrupted interaction with Ssa1. It is interesting to note that the 6KR mutant displayed a higher-than-normal Ssa1-Ydj1 interaction, which may suggest that wild-type Ydj1 is basally acetylated to some extent.

The current model of client processing is the client is first bound by Hsp40 and then delivered to Hsp70. This, in turn, activates the ATPase activity of Hsp70, and both Hsp70 and Hsp40 interact with the middle domain of Hsp90. Hsp70 and Hsp40 eventually dissociate from this transient complex, leaving the client with Hsp90 for the final stages of client maturation [10]. The driving forces behind client transfer from Hsp70 to Hsp90 remain obscure. It has been proposed in mammalian cells that the NudC protein is a key mediator of chaperone-client transfer, but the protein is not present in yeast [42]. Data from our MS and IP experiments suggest that Ydj1 acetylation disrupts the Ydj1-Ssa1 interaction while strengthening the Ydj1-Hsc82 interaction. Taken together, our data suggest that Ydj1 acetylation may be a key component of the protein folding process.

A large number of our Ydj1 interactors were associated with protein translation. Several previous studies have hinted at a connection between Ydj1 and protein synthesis. For example, the temperature-sensitive Ydj1 mutant *ydj1-151*, displays defects in the translation of two heterologously expressed proteins, firefly luciferase and GFP [43]. Additionally, cells lacking Ydj1 are sensitive to inhibitors of translation such as hygromycin B and cycloheximide [44]. Protein synthesis is carried out by the ribosome, an RNA-protein complex consisting of a small and large subunit [45]. Ribosome assembly involves more than 200 assembly factors, 79 ribosomal proteins, rRNAs, and other ribosomal-related proteins that participate in the complex pathway to achieve protein synthesis [45,46]. Assembly of the ribosome begins with transcription of rRNA in the nucleolus, then the rRNA undergoes complex folding, nucleotide modification, and binding to ribosomal proteins [46]. In *S. cerevisiae*, the small subunit (40S) has 33 ribosomal proteins and an 18S ribosomal RNA [47]. The large subunit (60S) has 46 ribosomal proteins and three ribosomal RNAs [45]. The small subunit is the location of the decoding site, where the anticodon of an amino-acyl tRNA base pairs with its respective codon found in the mRNA [45]. The large subunit contains the peptidyl transferase center (PTC), which is the active site of the unit. This is where rRNA catalyzes the formation of peptide bonds and hydrolyzes the peptidyl-tRNA bond [45]. Many of the interactors of Ydj1 identified in our screen were structural components of the ribosome, both from the small and large subunit. Recent studies have revealed that Hsp90 is primarily associated with the small subunit, while Hsp70 (which binds the nascent chain) was found to be associated with all the ribosome fractions [48]. Rps0, Rps3, and Gcd11 were not only identified as Ydj1 interactors, but their abundance was drastically reduced in cells lacking wild-type Ydj1 and expressing 6KQ. Overall, this suggests that these proteins are novel *bona fide* clients of Ydj1. Several PTMs, such as lysine ubiquitination, play an important role in the regulation of protein stability, such as the rise and fall of cyclins throughout the cell cycle. The K37Q mutant displays a higher abundance than wild-type Ydj1, and as our mutants were expressed from a constitutive promoter, this change in abundance is likely caused by altered protein degradation. We have considered the possibility that K37 may also be a site of ubiquitination. In this scenario, acetylation would prevent ubiquitination, leading to a stabilization of Ydj1; however, the levels of Ydj1 in the K37R mutant do not corroborate this theory. It is, of course, possible that K37Q prevents ubiquitination at a lysine not studied in this project, which will be investigated further in the future. Farnesylation of Ydj1, which can be detected as a doublet on Western Blots, can be clearly seen in all mutant samples. This suggests that while acetylation may impact association with Ssa1, farnesylation of Ydj1 remains unaffected.

Overall, our results suggest that acetylation of the residues located in the J-domain (particularly K23 and K37) disrupts Ssa1-Ydj1 interaction, thus inhibiting a large proportion of its cellular activity. Our data thus produce a conundrum: Why would the cell want to inhibit Ydj1 activity? The G1 cyclin Cln3 competes with Ydj1 for binding to Ssa1 during cell cycle progression and nutrient starvation [49]. Follow-up studies determined that phosphorylation of Ssa1 on T36 was required for this process, allowing displacement of Ydj1 from Ssa1 and allowing Cln3 to bind and be targeted for destruction [50]. It is thus possible that Ydj1 acetylation is part of the same mechanism to regulate Ssa1-Ydj1 interaction during cell cycle progression. Similarly, our lab has shown that deacetylation of Ssa1 is needed for a robust heat shock response [27]. This evidence, combined with our data, suggests that deacetylation of both Ssa1 and Ydj1 may be coordinated to increase Ssa1-Ydj1 interaction and trigger refolding of important client complexes under stressful conditions.

Taken together, this study provides evidence that the J-domain acetylation of Ydj1 has a significant impact on its chaperone and client interactions, leading to altered stress resistance. While beyond the scope of this study, we envisage future work that explores the conditions and enzymes that control Ydj1 acetylation. HDAC6 deacetylates the Ydj1 homolog DNAJA1 in mammalian cells, a process required for Hsp70-DNAJA1 interaction [23]. Although the specific sites of acetylation on DNAJA1 have not yet been identified, it is possible from the evidence presented in this study that K23 and K37 equivalent sites on DNAJA1 are responsible. Although these studies of the chaperone code are still in their infancy, it will be exciting to explore not only the role and regulation of chaperone and client PTMs but also their interplay in proteostasis.

## Experimental procedures

### Reagents and resources

Details on all reagents and resources (yeast strains and plasmids) are provided in Supplementary Table S1.

### Yeast Strains and growth conditions

Yeast cultures were grown in either YPD (1% yeast extract, 2% glucose, 2% peptone) or grown in SD (0.67% yeast nitrogen base without amino acids and carbohydrates, 2% glucose) supplemented with the appropriate nutrients to select for plasmids and tagged genes. *Escherichia coli* DH5α was used to propagate all plasmids. *E. coli* cells were cultured in Luria broth medium (1% Bacto tryptone, 0.5% Bacto yeast extract, 1% NaCl) and transformed to ampicillin or kanamycin resistance by standard methods. For monitoring stress conditions, cells were grown to mid-log phase, 10-fold serially diluted, and then plated onto appropriate media using a 48-pin replica-plating tool. Images of plates were taken after three days at 30 °C.

### Plasmid construction

The plasmid for expressing Ydj1 with a C-terminal FLAG tag (pYCP-GPD-YDJ1-FLAG) was constructed by VectorBuilder, and acetylation site mutants were generated by Genscript. Plasmids are listed in supplementary data (Table S1), and full plasmid sequences in Snapgene format are available on request. For bacterial expression, the *YDJ1* gene was PCR amplified from yeast genomic DNA to produce overhangs containing XbaI and XhoI restriction sites on the 5’ and 3’ ends of the PCR product. After digestion with XbaI and XhoI, the YDJ1-containing fragment was inserted into *E. coli* expression plasmid pRSETA (Thermo) resulting in a construct that allows expression of a 6xHIS-TEV-Ydj1 fusion protein.

### Expression of recombinant Ydj1 in *E. coli*

HIS-Ydj1 expression plasmids were transformed into *E. coli* BL21-competent cells. Cells were grown to an OD_600_=0.6 in 2YT media supplemented with 100 μg/ml ampicillin at 37 °C. Once the appropriate OD was reached, protein expression was induced by adding IPTG at a 1 mM final concentration. Cultures were incubated at 37 °C with shaking for an additional 4 hrs. Induced cells were then harvested via centrifugation (5000 rpm, 5 min) and sonicated in intervals of 30 seconds on ice until clear. Protein extracts were stored at −20 ℃ until prepared for Western Blot analysis.

### Immunoprecipitation

For FLAG IP, cells were harvested, and FLAG-tagged proteins were isolated as follows: Protein was extracted via bead beating in 500 μl binding buffer (50 mM Na-phosphate pH 8.0, 300 mM NaCl, 0.01% Tween-20). 200 μg of protein extract was incubated with 30 μl anti-Flag M2 magnetic beads (Sigma) at 4 C overnight. Anti-FLAG M2 beads were collected by the magnet and then washed 5 times with 500 μl binding buffer. After the final wash, the buffer was aspirated, and beads were incubated with 65 μl Elution buffer (binding buffer supplemented with 10 μg/ml 3X FLAG peptide (Apex Bio) for 1 hr at room temperature, then beads were collected via magnet. The supernatant containing purified FLAG-protein was transferred to a fresh tube, 25 μl of 4X SDS-PAGE sample buffer was added, and the sample was denatured for 5 min at 95 C. The eluates were separated by SDS-PAGE (7.5–10%) and visualized by immunoblotting.

### Immunoblotting

Protein extracts were made as described in [22]. 20 μg of protein was separated by 4%– 12% NuPAGE SDS-PAGE (Thermo Fisher Scientific). Proteins were detected using the primary antibodies incubated with primary antibodies (Supplementary Data 5) with the following dilutions: anti-GAPDH (1:5000), anti-HA-tag (1:2000), anti-FLAG tag (1:2000), anti-His tag (1:2000), anti-Ssa1 (1:2000), and anti-PGK1 (1:5000). Membranes were washed with TBS-Tween 20 (0.2%) and incubated with the corresponding secondary antibodies: anti-rat IgG-HRP (1:5000), anti-mouse IgG-HRP (1:5000), anti-rabbit IgG-HRP (1:5000), and anti-mouse IgM-HRP (1:5000). Blots were imaged on a ChemiDoc MP imaging system (Bio-Rad). After treatment with SuperSignal West Pico Chemiluminescent Substrate (Thermo Fisher Scientific). Blots were stripped and re-probed with the relevant antibodies using Restore Western Blot Stripping Buffer (Thermo Fisher Scientific).

### Proteins

Ssa1 was isolated as previously described [51]. His-tagged Ydj1 wild-type and mutant proteins were expressed in *E. coli* Rosetta (DE3) cells in the presence of IPTG. His-tagged Ydj1 proteins were purified using Ni-sepharose (Cytiva) in Ni buffer (40 mM Hepes (pH 7.5), 0.3 M NaCl, 10% glycerol(vol/vol)). Bound protein was washed with Ni buffer containing 10 mM imidazole followed by 25 mM imidazole and then eluted with Ni-buffer containing 250 mM imidazole. Ydj1 containing Ni-sepharose fractions were pooled and chromatographed on a HiLoad Superdex 200 size exclusion column in 25 mM Hepes (pH 7.5), 0.1 M NaCl, 0.1 mM EDTA, 1 mM DTT, 5% glycerol (vol/vol) using an AKTA pure 25 system. For BLI experiments, Yd1 proteins were labeled using a 1.5-fold excess of NHS-PEG4-Biotin (Thermo Scientific, Life Technologies). Excess biotin reagent was removed using 7K molecular weight cut-off Zeba Spin Desalting Columns (Thermo Scientific). Concentrations given are for Ydj1 dimers and for Ssa1 and luciferase monomers.

### Biolayer interferometry (BLI) analysis

BLI was used to monitor the interaction between Ssa1 and Ydj1 wild-type or mutant proteins using a Sartorius Octet R4 instrument and streptavidin biosensors at 23 °C. Wild-type Ydj1-biotin (5 μg/ml) was loaded on the biosensors and the association of Ssa1 (1 μM) with Ydj1 was monitored over time followed by dissociation in the absence of Ssa1. Interaction of Ssa1 with Ydj1 mutant proteins was determined using a loading of Ydj1-biotin mutant proteins equivalent to Ydj1-biotin wild-type. All Ydj1, Ssa1 BLI steps were performed in 25 mM Hepes pH 7.5, 50 mM KCl, 2 mM DTT, 5 mM MgCl_2_, 0.5 mM ATP and 0.02% Tween-20 (vol/vol). Ssa1 nonspecific binding was monitored using a reference biosensor subjected to each of the above steps in the absence of the biotinylated Ydj1, and the nonspecific binding signal was subtracted from the corresponding experiment.

### Luciferase reactivation assay

Luciferase (8 μM) was chemically denatured in 5 M guanidine hydrochloride, 25 mM Hepes pH 7.5, 50 mM KCl, 0.5 mM EDTA, for 10 min at 23 °C. For reactivation, denatured luciferase was diluted 100-fold into 25 mM Hepes pH 7.5, 0.1 M KOAc, 5 mM DTT, 10 mM Mg(OAc)_2_, 2 mM ATP, an ATP regenerating system (10 mM creatine phosphate, 3 μg creatine kinase), 3 μM Ssa1 and 0.3 μM Ydj1 wild-type or mutant. Aliquots were removed at the indicated times, and light output was measured using a Tecan Spark in luminescence mode with an integration time of 1000 ms following the injection of luciferin (50 μg/ml). Reactivation was determined compared to a non-denatured luciferase control.

### ATPase assay

Steady-state ATP hydrolysis was measured at 37 °C in 25 mM Hepes pH 7.5, 50 mM KCl, 2 mM DTT, 0.01 % (vol/vol) Triton X-100, 5 mM MgCl_2_, and 2 mM ATP using a pyruvate kinase/lactate dehydrogenase enzyme-coupled assay as described [52] and 1 μM Ssa1 and 1 μM Ydj1 wild-type or mutant.

## Mass Spectrometry

### In-solution sample digestion and desalting

Ydj1 complex immunoprecipitates were eluted from beads using 8M Urea,10mM DTT in 50 mM Tris pH 8.5 for 45 min at room temperature with mixing at 600 rpm before alkylation with 50 mM IAA for 30 min in the dark. Samples were diluted 6x with 50 mM Tris pH 8.5 to reach a 2 M Urea concentration and digested with 0.4 µg of trypsin-LysC mix (Promega) overnight at 37°C. Tryptic peptides were desalted with Pierce C18 Desalting Spin Columns (Thermo Fisher Scientific) according to the manufacturer’s protocol, dried down on SpeedVac, and resuspended in mobile phase A (0.2% formic acid in water) immediately prior to mass spectrometric analysis.

### Liquid chromatography-tandem mass spectrometry peptide analysis

Resuspended peptides were separated by nanoflow reversed-phase liquid chromatography (LC). An Ultimate 3000 UHPLC (Thermo Scientific) was used to load ∼1 μg of peptides on the column and separate them at a flow rate of 300 nL/min. The column was a 15 cm long EASY-Spray C18 (packed with 2 µm PepMap C18 particles, 75 µm i.d., Thermo Scientific). The analytical gradient was performed by increasing the relative concentration of mobile phase B (0.2% formic acid, 4.8% water in acetonitrile) in the following steps: from 2% to 30% in 32 min, from 30% to 50% in 5 min, and from 50 to 85% in 5 min (for washing the column). The 4 min wash at high organic concentration was followed by moving to 15% in 2 minutes, increasing to 70% in 1 min for a secondary wash before re-equilibration of the column at 2% B for 7.5 min, for a total run time of 68 min. A 2.2 kV potential was applied to the column outlet using an in-house nanoESI source based on the University of Washington design for generating nano-electrospray. All mass spectrometry (MS) measurements were performed on a tribrid Orbitrap Eclipse (Thermo Scientific). Broadband mass spectra (MS1) were recorded in the Orbitrap over a 375-1500 m/z window, using a resolving power of 120,000 (at 200 m/z) and an automatic gain control (AGC) target of 4e5 charges (maximum injection time: 50 ms). Precursor ions were quadrupole selected (isolation window: 1.6 m/z) based on a data-dependent logic, using a maximum duty cycle time of 3 s. Monoisotopic precursor selection and dynamic exclusion (60 s) were applied. Peptides were filtered by the intensity and charge state, allowing the fragmentation only of precursors from 2+ to 7+. Tandem mass spectrometry (MS2) was performed by fragmenting each precursor passing the selection criteria using both higher energy collisional dissociation (HCD) with normalized collision energy (NCE) set at 30% and electron transfer dissociation – higher energy collisional dissociation (EThcD), with ETD reagent target set at 5e5, reaction time calculated on the basis of a calibration curve and supplemental collisional activation set at NCE=10%. The AGC target for both HCD and EThcD MS2 was set at 8e4 (maximum injection time: 55 ms), and spectra were recorded at 15,000 resolving power.

### MS Data analysis

For general identification of all proteins included in the samples, HCD fragmentation data were processed with Protein Discoverer 2.4 utilizing Sequest HT and MS Amanda search engines. For both Precursor Mass Tolerance was 10 ppm and Fragment Mass Tolerance was 0.2 Da. Carbamidomethylation (C) was allowed as a static modification, and dynamic modifications were as follows: Oxidation(M), Acetyl (protein N-term), and Acetyl (K). Identified peptides were validated using Percolator, and the target FDR value was set to 0.01 (strict) and 0.05 (relaxed). Finally, results were filtered for high-confidence peptides using consensus steps. A control peptide error rate strategy was used, and 0.01 (strict) and 0.05 (relaxed) values were used for Target FDR for both PSM and Peptide levels. Changes in protein abundance between the two strains were statistically tested by ANOVA using the built-in function within Proteome Discoverer.

## Data Availability

The mass spectrometry proteomic data have been deposited to the ProteomeXchange Consortium (http://proteomecentral.proteomexchange.org) via the MassIVE partner repository with the dataset identifier MSV000094536 (Reviewer access at ftp://MSV000094536@massive.ucsd.edu; reviewer password: Ydj1_Acetylation) and cross-listed on PRIDE (PXD051428).

## Gene ontology enrichment analysis

Gene ontology analysis of Ydj1 immunoprecipitated complexes was accomplished using TheCellMap.org.

## General data analyses

Data processing and analyses were performed using GraphPad Prism (version 7).

## Acknowledgments

This work was supported by the NIH (R15GM139059, R21NS133682 and R01GM139885 to AWT, R35SGM147397 to LF), National Science Foundation Graduate Research Fellowship Program under Grant No. 2235055 (MMM), the Intramural Research Program of the NIH, NCI, Center for Cancer Research (SW). We acknowledge the PRIDE team for the deposition of our data to the ProteomeXchange consortium.

## Author Contributions

SO, CS, and N completed all experiments in Figures 2-5, JRH and SW completed experiments in Figures 6 and S1. JTK and LF completed experiments in Figure 4. AT acquired project funding and designed all the experiments for this project. MM analyzed the data for Figure 1. All authors contributed to the writing of the manuscript.

## References

1. Craig EA, Gambill BD, Nelson RJ. Heat shock proteins: molecular chaperones of protein biogenesis. Microbiol Rev. 1993;57: 402–414.

2. Kampinga HH, Craig EA. The HSP70 chaperone machinery: J proteins as drivers of functional specificity. Nat Rev Mol Cell Biol. 2010;11: 579–592.

3. Qiu X-B, Shao Y-M, Miao S, Wang L. The diversity of the DnaJ/Hsp40 family, the crucial partners for Hsp70 chaperones. Cell Mol Life Sci. 2006;63: 2560–2570.

4. Ruger-Herreros C, Svoboda L, Mogk A, Bukau B. Role of J-domain Proteins in Yeast Physiology and Protein Quality Control. J Mol Biol. 2024; 168484.

5. Fan C-Y, Ren H-Y, Lee P, Caplan AJ, Cyr DM. The type I Hsp40 zinc finger-like region is required for Hsp70 to capture non-native polypeptides from Ydj1. J Biol Chem. 2005;280: 695–702.

6. Caplan AJ, Douglas MG. Characterization of YDJ1: a yeast homologue of the bacterial dnaJ protein. J Cell Biol. 1991;114: 609–621.

7. Wright CM, Fewell SW, Sullivan ML, Pipas JM, Watkins SC, Brodsky JL. The Hsp40 molecular chaperone Ydj1p, along with the protein kinase C pathway, affects cell-wall integrity in the yeast Saccharomyces cerevisiae. Genetics. 2007;175: 1649–1664.

8. Millson SH, Truman AW, Piper PW. Hsp90 and phosphorylation of the Slt2(Mpk1) MAP kinase activation loop are essential for catalytic, but not non-catalytic, Slt2-mediated transcription in yeast. Cell Stress Chaperones. 2022;27: 295–304.

9. Mogk A, Bukau B, Kampinga HH. Cellular Handling of Protein Aggregates by Disaggregation Machines. Mol Cell. 2018;69: 214–226.

10. Wickramaratne AC, Liao J-Y, Doyle SM, Hoskins JR, Puller G, Scott ML, et al. J-domain Proteins form Binary Complexes with Hsp90 and Ternary Complexes with Hsp90 and Hsp70. J Mol Biol. 2023;435: 168184.

11. Kampinga HH, Andreasson C, Barducci A, Cheetham ME, Cyr D, Emanuelsson C, et al. Function, evolution, and structure of J-domain proteins. Cell Stress Chaperones. 2019;24: 7–15.

12. Jiang Y, Rossi P, Kalodimos CG. Structural basis for client recognition and activity of Hsp40 chaperones. Science. 2019;365: 1313–1319.

13. Tsai J, Douglas MG. A conserved HPD sequence of the J-domain is necessary for YDJ1 stimulation of Hsp70 ATPase activity at a site distinct from substrate binding. J Biol Chem. 1996;271: 9347–9354.

14. Sojourner SJ, Graham WM, Whitmore AM, Miles JS, Freeny D, Flores-Rozas H. The Role of HSP40 Conserved Motifs in the Response to Cytotoxic Stress. J Nat Sci. 2018;4. Available: https://www.ncbi.nlm.nih.gov/pubmed/29682607

15. Johnson JL, Craig EA. An essential role for the substrate-binding region of Hsp40s in Saccharomyces cerevisiae. J Cell Biol. 2001;152: 851–856.

16. Li J, Sha B. Structure-based mutagenesis studies of the peptide substrate binding fragment of type I heat-shock protein 40. Biochem J. 2005;386: 453–460.

17. Ayala Mariscal SM, Kirstein J. J-domain proteins interaction with neurodegenerative disease-related proteins. Exp Cell Res. 2021;399: 112491.

18. Velasco-Carneros L, Cuéllar J, Dublang L, Santiago C, Maréchal J-D, Martín-Benito J, et al. The self-association equilibrium of DNAJA2 regulates its interaction with unfolded substrate proteins and with Hsc70. Nat Commun. 2023;14: 5436.

19. Mitchem MM, Shrader C, Abedi E, Truman AW. Novel insights into the post-translational modifications of Ydj1/DNAJA1 co-chaperones. Cell Stress Chaperones. 2023;29: 1–9.

20. Hildebrandt ER, Cheng M, Zhao P, Kim JH, Wells L, Schmidt WK. A shunt pathway limits the CaaX processing of Hsp40 Ydj1p and regulates Ydj1p-dependent phenotypes. Elife. 2016;5. doi:10.7554/eLife.15899

21. Flom GA, Lemieszek M, Fortunato EA, Johnson JL. Farnesylation of Ydj1 is required for in vivo interaction with Hsp90 client proteins. Mol Biol Cell. 2008;19: 5249–5258.

22. Sluder IT, Nitika, Knighton LE, Truman AW. The Hsp70 co-chaperone Ydj1/HDJ2 regulates ribonucleotide reductase activity. PLoS Genet. 2018;14: e1007462.

23. Zhang L, Liu S, Liu N, Zhang Y, Liu M, Li D, et al. Proteomic identification and functional characterization of MYH9, Hsc70, and DNAJA1 as novel substrates of HDAC6 deacetylase activity. Protein Cell. 2015;6: 42–54.

24. Scroggins BT, Robzyk K, Wang D, Marcu MG, Tsutsumi S, Beebe K, et al. An acetylation site in the middle domain of Hsp90 regulates chaperone function. Mol Cell. 2007;25: 151– 159.

25. Seo JH, Park J-H, Lee EJ, Vo TTL, Choi H, Kim JY, et al. ARD1-mediated Hsp70 acetylation balances stress-induced protein refolding and degradation. Nat Commun. 2016;7: 12882.

26. Krämer OH, Mahboobi S, Sellmer A. Drugging the HDAC6-HSP90 interplay in malignant cells. Trends Pharmacol Sci. 2014;35: 501–509.

27. Xu L, Nitika, Hasin N, Cuskelly DD, Wolfgeher D, Doyle S, et al. Rapid deacetylation of yeast Hsp70 mediates the cellular response to heat stress. Sci Rep. 2019;9: 16260.

28. Aksnes H, Drazic A, Marie M, Arnesen T. First Things First: Vital Protein Marks by N-Terminal Acetyltransferases. Trends Biochem Sci. 2016;41: 746–760.

29. Caplan AJ, Tsai J, Casey PJ, Douglas MG. Farnesylation of YDJ1p is required for function at elevated growth temperatures in Saccharomyces cerevisiae. J Biol Chem. 1992;267: 18890–18895.

30. Nitika, Porter CM, Truman AW, Truttmann MC. Post-translational modifications of Hsp70 family proteins: Expanding the chaperone code. J Biol Chem. 2020;295: 10689–10708.

31. Rosenzweig R, Nillegoda NB, Mayer MP, Bukau B. The Hsp70 chaperone network. Nat Rev Mol Cell Biol. 2019;20: 665–680.

32. Alao JP. The regulation of cyclin D1 degradation: roles in cancer development and the potential for therapeutic invention. Mol Cancer. 2007;6: 24.

33. Gaur D, Singh P, Guleria J, Gupta A, Kaur S, Sharma D. The Yeast Hsp70 Cochaperone Ydj1 Regulates Functional Distinction of Ssa Hsp70s in the Hsp90 Chaperoning Pathway. Genetics. 2020;215: 683–698.

34. Bukau B, Deuerling E, Pfund C, Craig EA. Getting newly synthesized proteins into shape. Cell. 2000;101: 119–122.

35. Ben-Shem A, Jenner L, Yusupova G, Yusupov M. Crystal structure of the eukaryotic ribosome. Science. 2010;330: 1203–1209.

36. Takacs JE, Neary TB, Ingolia NT, Saini AK, Martin-Marcos P, Pelletier J, et al. Identification of compounds that decrease the fidelity of start codon recognition by the eukaryotic translational machinery. RNA. 2011;17: 439–452.

37. Nitika, Truman AW. Cracking the Chaperone Code: Cellular Roles for Hsp70 Phosphorylation. Trends Biochem Sci. 2017;42: 932–935.

38. Nitika, Zheng B, Ruan L, Kline JT, Omkar S, Sikora J, et al. Comprehensive characterization of the Hsp70 interactome reveals novel client proteins and interactions mediated by posttranslational modifications. PLoS Biol. 2022;20: e3001839.

39. Woodford MR, Backe SJ, Wengert LA, Dunn DM, Bourboulia D, Mollapour M. Hsp90 chaperone code and the tumor suppressor VHL cooperatively regulate the mitotic checkpoint. Cell Stress Chaperones. 2021;26: 965–971.

40. Kaluarachchi Duffy S, Friesen H, Baryshnikova A, Lambert J-P, Chong YT, Figeys D, et al. Exploring the yeast acetylome using functional genomics. Cell. 2012;149: 936–948.

41. Lin Y-Y, Qi Y, Lu J-Y, Pan X, Yuan DS, Zhao Y, et al. A comprehensive synthetic genetic interaction network governing yeast histone acetylation and deacetylation. Genes Dev. 2008;22: 2062–2074.

42. Biebl MM, Delhommel F, Faust O, Zak KM, Agam G, Guo X, et al. NudC guides client transfer between the Hsp40/70 and Hsp90 chaperone systems. Mol Cell. 2022;82: 555– 569.e7.

43. Brodsky JL, Lawrence JG, Caplan AJ. Mutations in the cytosolic DnaJ homologue, YDJ1, delay and compromise the efficient translation of heterologous proteins in yeast. Biochemistry. 1998;37: 18045–18055.

44. Gillies AT, Taylor R, Gestwicki JE. Synthetic lethal interactions in yeast reveal functional roles of J protein co-chaperones. Mol Biosyst. 2012;8: 2901–2908.

45. Klinge S, Woolford JL Jr. Ribosome assembly coming into focus. Nat Rev Mol Cell Biol. 2019;20: 116–131.

46. Woolford JL Jr, Baserga SJ. Ribosome biogenesis in the yeast Saccharomyces cerevisiae. Genetics. 2013;195: 643–681.

47. Moraleva AA, Deryabin AS, Rubtsov YP, Rubtsova MP, Dontsova OA. Eukaryotic Ribosome Biogenesis: The 40S Subunit. Acta Naturae. 2022;14: 14–30.

48. Kolhe JA, Babu NL, Freeman BC. The Hsp90 molecular chaperone governs client proteins by targeting intrinsically disordered regions. Mol Cell. 2023;83: 2035–2044.e7.

49. Vergés E, Colomina N, Garí E, Gallego C, Aldea M. Cyclin Cln3 is retained at the ER and released by the J chaperone Ydj1 in late G1 to trigger cell cycle entry. Mol Cell. 2007;26: 649–662.

50. Truman AW, Kristjansdottir K, Wolfgeher D, Hasin N, Polier S, Zhang H, et al. CDK-dependent Hsp70 Phosphorylation controls G1 cyclin abundance and cell-cycle progression. Cell. 2012;151: 1308–1318.

51. Reidy M, Sharma R, Roberts B-L, Masison DC. Human J-protein DnaJB6b Cures a Subset of Saccharomyces cerevisiae Prions and Selectively Blocks Assembly of Structurally Related Amyloids. J Biol Chem. 2016;291: 4035–4047.

52. Tamura JK, Gellert M. Characterization of the ATP binding site on Escherichia coli DNA gyrase. Affinity labeling of Lys-103 and Lys-110 of the B subunit by pyridoxal 5’-diphospho-5’-adenosine. J Biol Chem. 1990;265: 21342–21349.

